# The DNA damage response pathway is required for multiciliated cell differentiation

**DOI:** 10.1101/2025.07.13.664567

**Authors:** Cayla E Jewett, Andrew J Holland, Chad G Pearson

## Abstract

DNA damage can result from external sources or occur during programmed genome rearrangements in processes like immunity or meiosis. To maintain genome integrity, cells activate DNA repair pathways that prevent harmful outcomes such as cancer or immune dysfunction. In this study, we uncover a novel role for DNA damage during the terminal differentiation of multiciliated cells (MCCs). MCCs, which line the airways, reproductive tracts, and brain ventricles, produce hundreds of motile cilia, each anchored by a centriole. Therefore, MCCs must generate hundreds of centrioles during differentiation. Normally, centriole duplication is tightly linked to the S and G2 phases of the cell cycle, raising questions about how MCCs override numerical and temporal restrictions on centriole duplication. We find that differentiating MCCs accumulate extensive double-stranded DNA breaks during centriole amplification, with damage levels correlating with centriole number. DNA damage response (DDR) kinases are essential for supporting centriole biogenesis and ciliogenesis. We also observe that transcriptional activity, required for the expression of centriole and cilia genes, produces RNA-DNA hybrids (R-loops) that co-localize with DNA damage. This suggests that transcription-coupled DNA damage helps initiate a pseudo–cell cycle program, permitting centriole amplification without triggering full S/G2 phase processes. Our findings indicate that MCCs harness DDR signaling as part of their developmental program, revealing a broader principle in which the canonical cell cycle is adaptively rewired during differentiation.

## Results

### DNA damage occurs during centriole amplification in MCCs

To address how MCCs undergo a unique cell cycle allowing for overduplication of centrioles in the absence of other S and G2 process, we screened for events that regulate the cell cycle. Surprisingly, differentiating cells that express the MCC transcription factor FOXJ1 produce abundant double-stranded DNA breaks labeled by γH2AX stained foci (Figure 1A, cyan nuclei label MCCs). These DNA breaks were not present in neighboring stem cells (Figure 1A). γH2AX foci in MCCs colocalized with the DNA damage marker 53BP1 (Figure 1B), suggesting that DNA repair machinery is recruited to damage sites. To confirm the DNA damage observed with immunofluorescence, an alkaline comet assay was performed and showed that differentiating MCCs have an elongated comet tail when compared to the comet head (Figure 1C,D, S1A). In contrast, this comet tail was not present in undifferentiated stem cells. Together this shows that DNA damage occurs during MCC differentiation.

**Figure 1.**
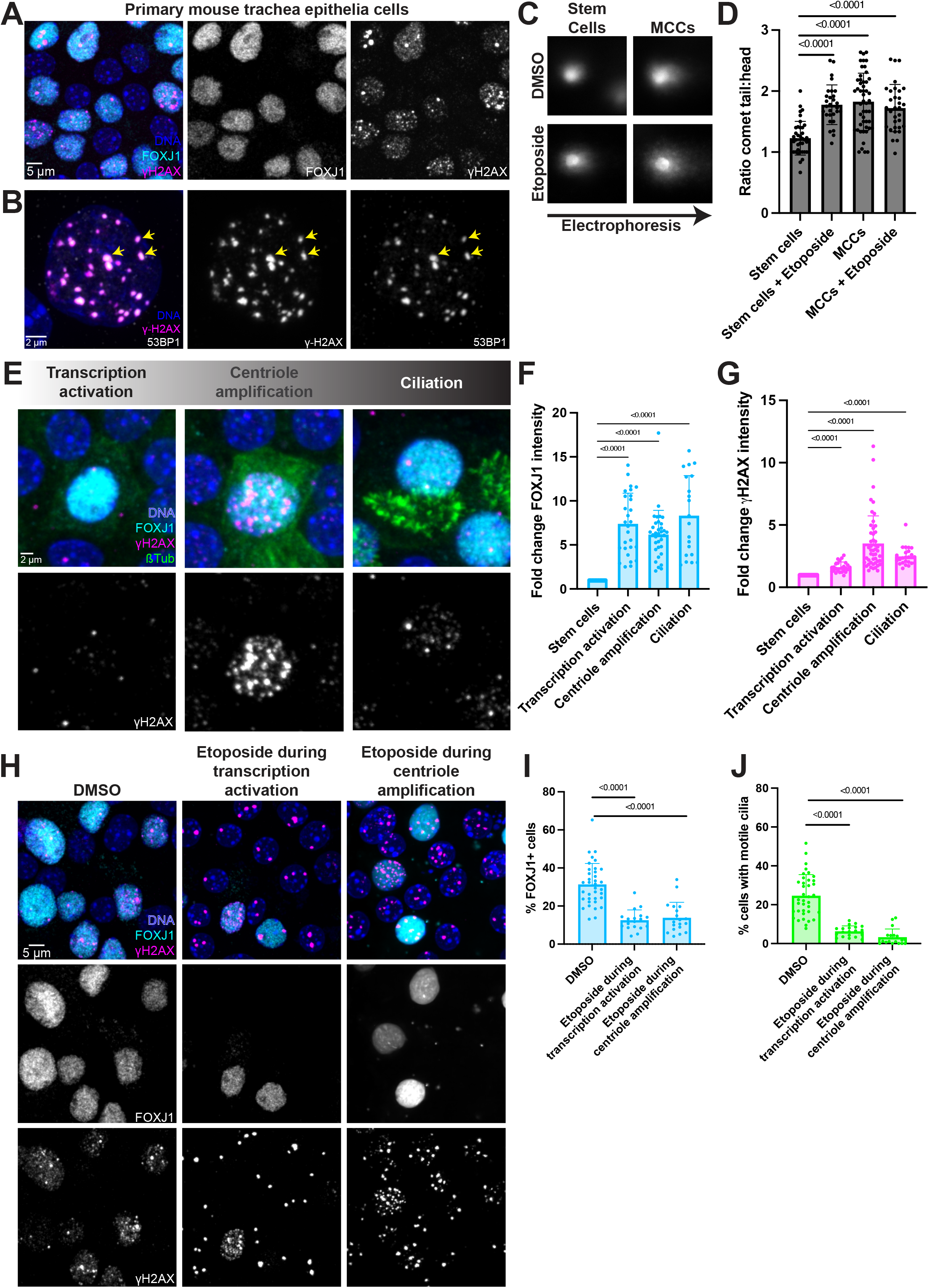
DNA damage occurs during centriole amplification in MCCs. (A) Primary mouse trachea epithelia cells at differentiation day 7, fixed, and stained for the MCC transcription factor (FOXJ1, cyan), DNA damage (γH2AX, magenta), and DAPI to mark nuclei. (B) Single nuclei of a MCC from primary mouse trachea epithelia cells cultured at an air-liquid interface for 7 days, fixed, and stained for DNA damage (γH2AX, magenta), DNA damage signaling (53BP1, gray), and DAPI to mark nuclei. Arrows point to colocalization of γH2AX and 53BP1 foci. (C) DMSO treated (top panel) and Etoposide treated (bottom panel) nuclei from comet assay showing stem cells prior to differentiation (left) and MCCs at differentiation day 5 (right). Comet tails indicating DNA breaks form in the electrophoresis direction. Note that MCCs at differentiation day 5 include both MCCs and stem cells without DNA breaks, contributing to the large spread of data. (D) Quantitation of comet assay measuring tail to head ratio. Nuclei with DNA breaks will have a longer comet tail. (E) Primary mouse trachea epithelia cells at differentiation days 3-14, fixed, and stained for the MCC transcription factor (FOXJ1, cyan), DNA damage (γH2AX, magenta), microtubules, centrioles, and cilia (β-tubulin, green), and DAPI to mark nuclei. MCCs were grouped into differentiation stages based on FOXJ1 and β-tubulin staining. (F-G) Fold-change in FOXJ1 (F) and γH2AX (G) nuclear intensities compared to neighboring stem cells. (H) Primary mouse trachea epithelia cells at differentiation day 7, fixed, and stained for the MCC transcription factor (FOXJ1, cyan), DNA damage (γH2AX, magenta), and DAPI to mark nuclei. Cells were treated with etoposide either during transcription activation (differentiation days 0-2) or during centriole amplification (differentiation days 3-5). (I-J) Percent of FOXJ1+ cells (I) and cells with motile cilia (J) for etoposide treatment in (H). All graphs show mean ± SD.

MCC differentiation occurs over two weeks in culture and can be asynchronous among a population of cells. Broadly, differentiation can be grouped into three stages. The first stage begins in the nucleus with transcription activation of a MCC-specific program. The second stage is cytoplasmic centriole amplification, when cells assemble hundreds of centrioles. The third stage is ciliation, during which cells nucleate motile cilia from these newly assembled centrioles. To address when during this process of differentiation DNA damage occurs, we analyzed γH2AX nuclear intensity alongside markers for the stages of MCC differentiation (Figure 1E, S1B). γH2AX intensity increased slightly during transcription activation, peaked at 4-fold during centriole amplification, and persisted throughout ciliation, although decreasing in intensity (Figure 1E-G). This timing of DNA damage during centriole amplification was consistent in both cultured primary mouse trachea epithelia cells and mouse tracheas *in vivo* (Figure 1E and S1C).

Together, this indicates that DNA damage occurs during the centriole amplification stage of differentiation.

To understand the precise timing of DNA damage, we used the auxin inducible degron (AID) system to degrade the earliest MCC transcription factor, GEMC1. GEMC1 is upstream of FOXJ1 and therefore GEMC1 loss completely prevents transcription activation and differentiation of MCCs^1-3^. To temporally control GEMC1 function, we knocked in an AID-Ruby tag at the endogenous GEMC1 locus in mice expressing the E3 ligase adaptor OsTIR1. GEMC1 depletion with IAA addition from the onset of differentiation resulted in loss of MCCs (Figure S1D), which is consistent with GEMC1 KO models^1-3^. Moreover, GEMC1 depletion abolished DNA damage (Figure S1D), suggesting that DNA damage is downstream of transcription activation and specific to MCC differentiation.

Given this robust physiologic DNA damage during MCC differentiation, we tested whether exogenous damage could induce differentiation by treating cells with the DNA damaging drug etoposide, either during the transcription activation stage or centriole amplification stage. Etoposide treatment during either stage did not induce differentiation and instead decreased the number of MCCs (Figure 1H-J). Furthermore, etoposide treatment generated γH2AX foci that were larger and fewer than those observed in MCCs (Figure 1H). Together this demonstrates that MCCs show physiologic DNA damage during differentiation that is unique from exogenous damage and correlates with centriole amplification.

### DNA damage scales with centriole number

Given that DNA damage correlates with the timing of centriole amplification, we then asked whether the amount of DNA damage relates to centriole number. Different multiciliated tissues have varying numbers of cilia and thus centrioles so we compared the amount of DNA damage across these MCC types relative to stem cells. This analysis revealed that nuclear γH2AX intensity was strongest in trachea MCCs (average 250-300 centrioles per cell^4,5^). Brain ventricle MCCs, which make an average of 40-100 centrioles^5-8^, still showed DNA damage during centriole amplification, but it was decreased relative to trachea MCCs (Figure 1E, 2A,B). Therefore, at the tissue level, MCCs with more centrioles have more DNA damage.

To further test the idea that DNA damage scales with the amount of centriole amplification, the quantity of DNA damage and centriole protein levels were assessed in individual trachea MCCs. The cytoplasmic protein intensity of four different centriole markers was quantified relative to nuclear γH2AX intensity. Centriole protein intensity positively correlated with γH2AX intensity (Figure 2C-E, S2A,B). In summary, DNA damage increases with centriole number in both tissues and individual MCCs.

**Figure 2.**
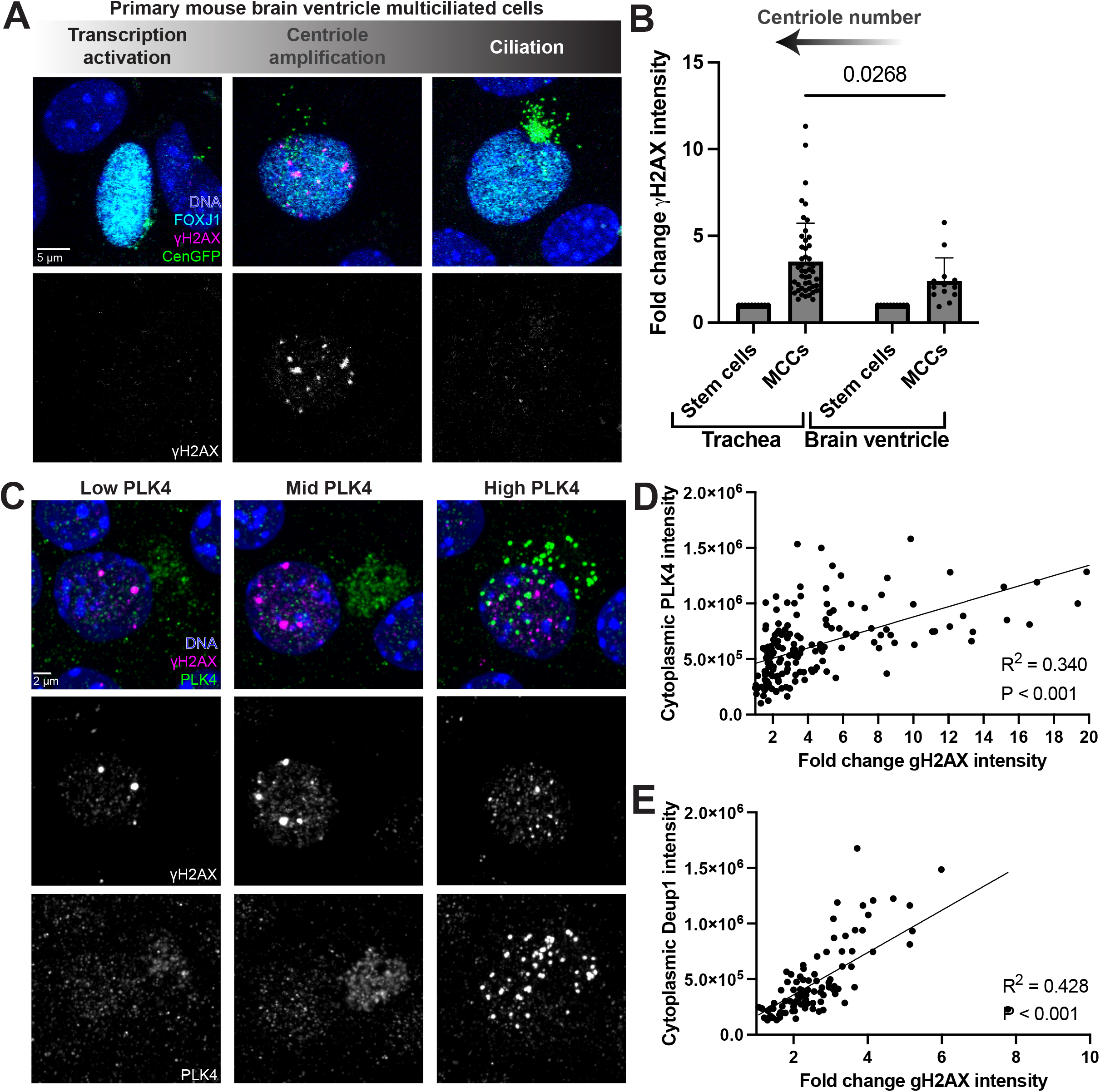
DNA damage scales with centriole number. (A) Primary mouse brain ventricle ependymal cells at differentiation day 7-10, fixed, and stained for the MCC transcription factor (FOXJ1, cyan), DNA damage (γH2AX, magenta), centrioles (CenGFP, green), and DAPI to mark nuclei. MCCs were grouped into differentiation stages based on FOXJ1 and CenGFP staining. (B) Fold-change in γH2AX nuclear intensities for trachea and brain ventricle MCCs compared to neighboring stem cells. Note that trachea MCCs have more centrioles than brain ventricle MCCs. (C) Primary mouse trachea epithelia cells at differentiation day 5, fixed, and stained for the centriole protein (PLK4, green), DNA damage (γH2AX, magenta), and DAPI to mark nuclei. (D-E) Linear correlation plot of cytoplasmic centriole protein intensity (PLK4 in (D) or Deup1 in (E)) versus fold-change in γH2AX nuclear intensity. Each dot represents a single cell. All graphs show mean ± SD.

### DNA damage sites contain repair machinery and RNA-DNA hybrids

DNA damage is a normally harmful process that cells rapidly repair. Our data demonstrate that MCCs show physiologic DNA damage, so we next explored the composition of these DNA damage sites. We first examined the localization of the three main DDR kinases ATR, ATM, and DNAPK, which are activated by phosphorylation^9^. We used phospho-antibodies to indicate the activity of ATR, ATM, DNAPK, or the substrates of DDR kinases, and analyzed nuclear intensity at the single cell level. pATM and pDNAPK showed a 3-fold intensity increase in cells undergoing centriole amplification compared to stem cells (Figure 3A-D, S3A). An antibody recognizing the phospho-motif of the DDR substrates also increased during centriole amplification and remained slightly elevated during the ciliation stage (Figure S1A). Moreover, foci of pATM, pDNAPK, and phosphorylated DDR substrates colocalized with γH2AX foci (Figure S3D-F). In contrast, pATR was only mildly increased in MCCs and did not colocalize with γH2AX foci (Figure S3G,H). Together this suggests that DNA damage sites are recruiting specific DDR machinery for signaling and repair.

**Figure 3.**
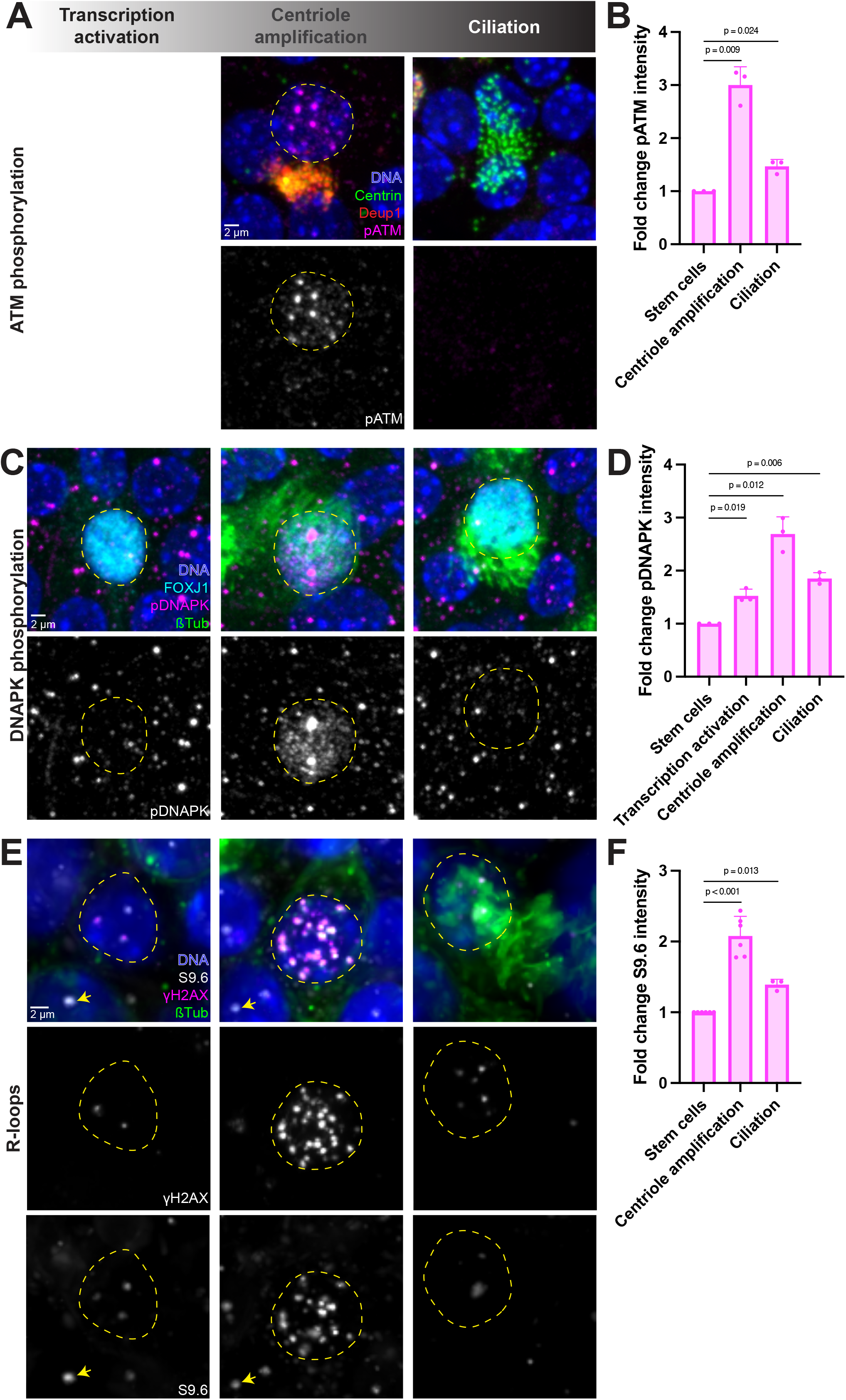
DNA damage sites contain repair machinery and RNA-DNA hybrids. (A) Primary mouse trachea epithelia cells at differentiation day 5, fixed, and stained for centrioles (Centrin, green), deuterosomes (Deup1, red), phosphorylated-ATM (pATM, magenta) and DAPI to mark nuclei. MCCs were grouped into differentiation stages based on Centrin and Deup1 staining, although due to antibody conflicts the transcription activation stage was unable to be identified with pATM staining. (B) Fold-change in pATM nuclear intensity compared to neighboring stem cells. (C) Primary mouse trachea epithelia cells at differentiation day 5, fixed, and stained for the MCC transcription factor (FOXJ1, cyan), phosphorylated-DNAPK (pDNAPK, magenta), microtubules, centrioles, and cilia (β-tubulin, green), and DAPI to mark nuclei. MCCs were grouped into differentiation stages based on FOXJ1 and β-tubulin staining. (D) Fold-change in pDNAPK nuclear intensity compared to neighboring stem cells. (E) Primary mouse trachea epithelia cells at differentiation day 5, fixed, and stained for the RNA-DNA hybrid/R-loop marker (S9.6, gray), DNA damage (γH2AX, magenta), microtubules, centrioles, and cilia (β-tubulin, green), and DAPI to mark nuclei. MCCs were grouped into differentiation stages based on γH2AX and β-tubulin staining. (F) Fold-change in S9.6 nuclear intensity compared to neighboring stem cells. All graphs show mean ± SD.

Some common sources of DNA damage include replication stress, transcription stress, and oxidative stress. The pseudo cell cycle state of MCCs permits centriole duplication without DNA replication^10^, however, replication stress can occur at telomeres in brain MCC progenitors with GEMC1 gain of function^7^. Therefore, we first tested whether the DNA damage observed during centriole amplification was at telomeres. DNA FISH with a telomere probe combined with γH2AX immunofluorescence revealed that most DNA damage sites were distinct from telomeres (Figure S3I,J), suggesting that telomeric replication stress is not the primary cause of DNA damage in differentiating trachea MCCs.

We next explored whether the unique transcriptome of MCCs requiring robust expression of many centriole and cilia genes could create a transcriptional burden contributing to DNA damage. R-loops are RNA-DNA structures that form in regions of heavy transcription and occur when a nascent RNA hybridizes to its DNA template displacing the non-template DNA strand^11^. R-loops are typically transient structures but if left unresolved cause DNA damage^11^. We therefore asked whether R-loops were present during MCC differentiation. To test this, MCCs were stained with the S9.6 antibody which recognizes R-loops^12^. In neighboring stem cells, R-loops could be visualized exclusively in the nucleolus (Figure 3E, arrows); however, in MCCs many R-loops were also present throughout the nucleus (Figure 3E, dashed circles). These R-loops formed specifically during the centriole amplification stage of MCC differentiation and colocalized with γH2AX foci (Figure 3E,F). Together, this suggests that during MCC differentiation, DNA damage sites recruit active repair kinases and contain DNA-RNA hybrid structures.

### DNA damage response kinases are required for centriole amplification

During a canonical cell cycle, DNA damage triggers the DDR pathway during which DDR kinases phosphorylate substrates ultimately activating a checkpoint that allows time for DNA repair before the cell cycle proceeds. Since MCCs undergo a pseudo cell cycle to amplify centrioles^10,13^, and these kinases are active during centriole amplification, we asked whether the activity of these kinases was required for MCC differentiation. To address this, kinase inhibitors to ATR, ATM, or DNAPK were used. Like many epithelial cell types, MCCs insert drug efflux pumps into the cell surface which can impact the effective drug concentration, especially for kinase inhibitors^14^. To overcome this, all drug experiments were performed in the presence of verapamil, a drug pump inhibitor. Also, kinase inhibitor concentrations were optimized with cell viability assays in proliferating mouse epithelial cells, dose response curves in MCCs, and validation with phospho-antibody staining (Figure S4A-F) (see methods).

We treated MCCs from the onset of differentiation with inhibitors to ATR, ATM, or DNAPK and examined the number of multiciliated cells one week later. Treatment with any of the kinase inhibitors reduced the number of MCCs (Figure 4A,B) suggesting that the DNA damage response pathway is required for MCC differentiation. We observed a slight decrease in the total number of cells under inhibitor treatment (Figure S4G). However, this was not enough to account for the decrease in MCCs, ruling out cell death as the cause for MCC loss. ATM and DNAPK inhibition had the most significant decrease in the number of MCCs (Figure 4A-B), and the largest increase in kinase activity during differentiation (Figure 3A-D).

**Figure 4.**
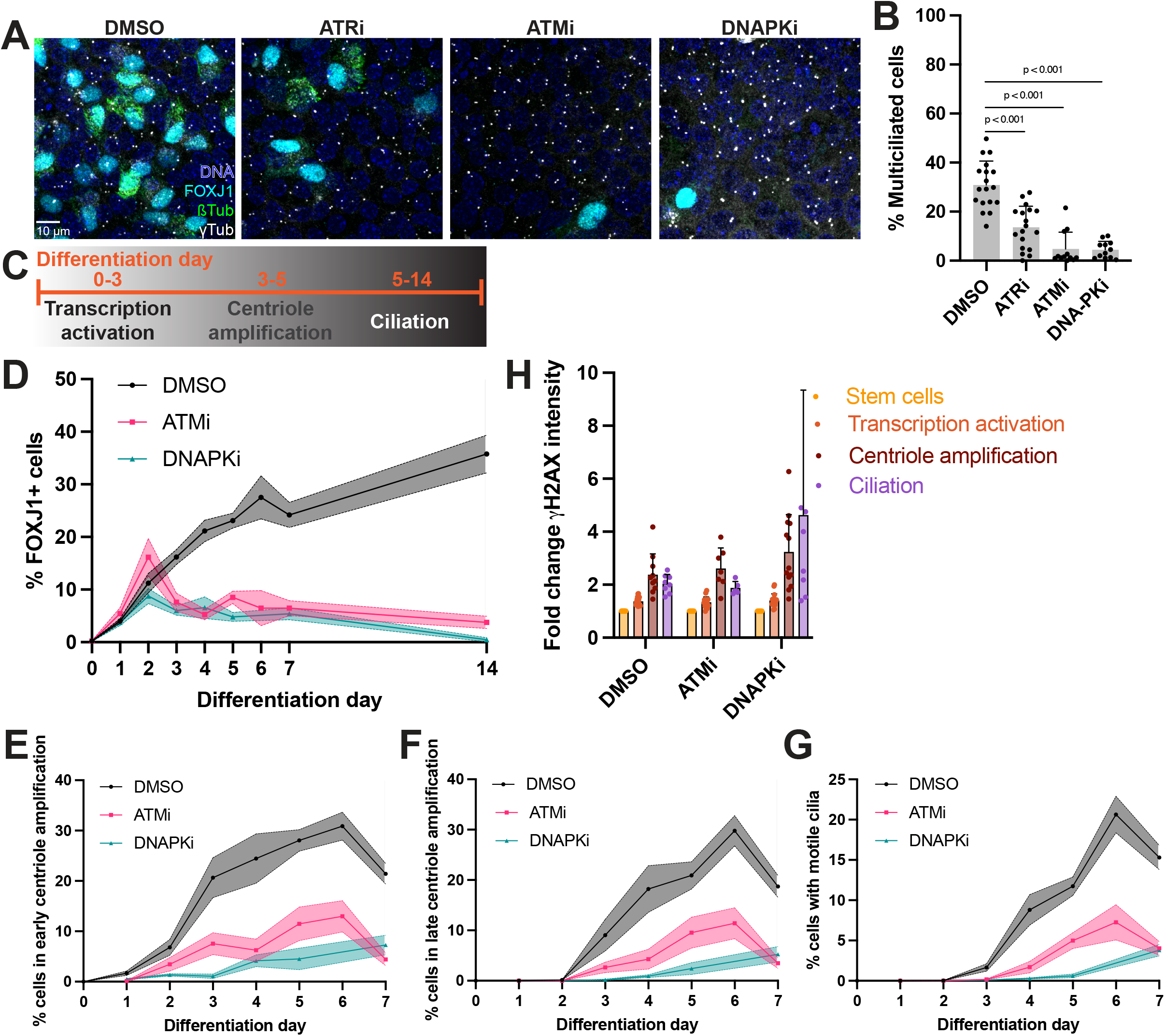
DNA damage response kinases are required for centriole amplification. (A) Primary mouse trachea epithelia cells at differentiation day 7, fixed, and stained for the MCC transcription factor (FOXJ1, cyan), centrioles (γTub, gray), microtubules, centrioles, and cilia (β-tubulin, green), and DAPI to mark nuclei. Cells were treated with DMSO plus verapamil or kinase inhibitors plus verapamil from the onset of differentiation to fixation (differentiation days 0-7). (B) Percent of FOXJ1+ multiciliated cells with kinase inhibitor treatment. (C) Schematic broadly depicting differentiation day timing in culture relative to stages of MCC differentiation. (D) Quantitation of time course experiments where primary mouse trachea epithelia cells were treated with DMSO plus verapamil or kinase inhibitors plus verapamil from the onset of differentiation and fixed every day for 7 days and again at 14 days. Graph shows the percent of FOXJ1+ cells throughout the time course. (E-G) Quantitation of time course experiments where primary mouse trachea epithelia cells were treated with DMSO plus verapamil or kinase inhibitors plus verapamil from the onset of differentiation and fixed every day for 7 days. Graphs show the percent of cells in early centriole amplification (E), late centriole amplification (F), and ciliation (G) stages. PCNT, CEP164, and β-tubulin were used to stage cells. (H) Fold-change in γH2AX nuclear intensity compared to neighboring stem cells for DMSO and kinase inhibitor treated cells at each stage of differentiation. B and H show mean ± SD. D-G show mean ± SEM.

Next, to temporally resolve kinase function throughout the stages of differentiation, cells were treated with kinase inhibitor from the onset of differentiation and fixed cells every day for one week and then at two weeks. ATM and DNAPK activity were not required at early timepoints (days 0-2) for transcription activation but were required for centriole amplification and ciliation (days 3-14) (Figure 4C, D). We next examined what happened to centrioles and cilia with kinase inhibition. ATM or DNAPK inhibition greatly delayed and reduced the number of cells in early centriole amplification, late centriole amplification, and ciliation stages (Figure 4D-G). This indicates that ATM and DNAPK activity are dispensable for transcription activation but required for centriole amplification and ciliation.

Finally, we asked whether DDR kinase activity was required for DNA damage in MCCs. Examining the initial γH2AX intensity profile revealed no change in γH2AX intensity during transcription activation, consistent with the timing of kinase function (Figure 4H). The few cells that did undergo centriole amplification or ciliation with kinase inhibition showed nuclear γH2AX at similar levels to control cells (Figure 4H, S4H). However, these centrioles were abnormal, and cilia were short, sparse, and disorganized compared to controls (Figure S4H,I), suggesting that compensatory DDR pathways may promote the MCC program but fail to assemble normal centrioles and cilia. Altogether, this demonstrates that DNA damage and the DNA damage response kinases are required for the centriole amplification and ciliation stages of MCC differentiation.

## Discussion

In trying to understand how MCCs repurpose the cell cycle to permit centriole overduplication, we discovered that MCCs show DNA damage during centriole amplification that scales with centriole number. In brain MCCs, DNA damage has been observed with overexpression of the earliest MCC transcription factor, GEMC1, and GEMC1 gain of function results in more centrioles^7^. The molecular pathways linking centriole copy number to DNA damage quantity remain an open area of investigation.

We found that DDR kinase activity is required for differentiation, but the purpose of DDR kinases could be multifaceted. Kinases may be required to repair DNA damage and prevent genomic instability. Alternatively, but not mutually exclusive, the kinases could activate a signaling pathway that triggers a checkpoint holding cells in a pseudo S-phase that allows time for centriole amplification. In brain MCCs, the p53/p21 pathway is activated in cycling progenitors that exit the cell cycle to differentiate^7^. Moreover, p21 is highly expressed in trachea MCCs during centriole amplification^10^. Given that DNA damage in canonical cycling cells delays the cell cycle until damage is repaired, it is possible that MCCs repurpose the DNA damage pathway to reprogram the cell cycle.

Our observation of RNA-DNA hybrids at damage sites suggests MCCs experience transcription stress that coincides with DNA damage. Further studies are needed to confirm this and rule out other sources of DNA damage. Replication stress is a common cause of DNA damage and given that centriole amplification is typically coupled to DNA replication, the unique cell cycle state of MCCs could lend itself to replication-induced damage. EdU incorporation to monitor DNA replication has been observed in MCCs when their pseudo cell cycle is perturbed^7,10,13,15^. However, EdU incorporation is not observed under normal MCC differentiation, and our data show that DNA damage is not at telomeres arguing against replication-associated damage. DNA damage has also been observed during massive genome remodeling events, such as the maternal to zygotic transition or neuronal activation^16-19^. Because MCCs must rewire their genome for robust expression of centriole and cilia genes, genome remodeling could contribute to DNA damage during differentiation. Moreover, ciliary beating requires ATP, so mitochondria activity is increased in MCCs to accommodate this increased ATP requirement^20-22^. While MCCs utilize mechanisms to reduce reactive oxygen species production^20^, the cellular energy demands could generate an oxidative stress causing DNA damage. Finally, MCCs line an epithelial surface where they are exposed to environmental insults. Therefore, a combination of factors could contribute to the DNA damage observed during MCC differentiation.

Identifying where DNA damage occurs within the genome will be an important clue to understanding the source of the damage. If DNA damage is from transcription, DNA damage should occur at or near heavily transcribed genes, such as centriole or cilia genes. We also observe heterogeneity in γH2AX foci size throughout the course of differentiation. It will be interesting to determine if different foci correlate with different levels of transcription, different genomic regions, or different sources of DNA damage.

Altogether our work identifies DNA damage as a prerequisite for centriole amplification in MCCs. Identifying how massive DNA damage is not harmful to MCCs and instead biologically programmed into the differentiation trajectory will illuminate how specialized cells maintain control over potentially pathological processes.

## Resource availability

Reagents generated in this study will be made available on request.

## Acknowledgments

We are grateful to Carolyn Ott and members of the Holland and Pearson labs for helpful discussions. We thank Margaret Strong and Craig Zikan for help with mouse husbandry. This work was funded by the Damon Runyon Cancer Research Foundation (CEJ is a Merck Fellow DRG-2478-22); a Hartwell Foundation Fellowship to CEJ; R01GM133897, R01GM114119, R01CA266199 to AJH; R35GM140813 to CGP.

## Author contributions

Conceptualization, CEJ, AJH, CGP; Investigation, CEJ; Formal Analysis, CEJ; Writing – Original Draft, CEJ, CGP; Writing –Review & Editing, CEJ, AJH, CGP; Funding Acquisition, CEJ, AJH, CGP.

## Materials and Methods

### Mouse Husbandry

Mice were housed and cared for in AAALAC-accredited facilities. All animal experiments were performed according to the Johns Hopkins University Institute Animal Care and Use Committee (MO21M300) or the University of Colorado Institute Animal Care and Use Committee (01490). Strains were maintained on a standard chow diet and a mixture of male and female mice were used for experiments. No differences in sexes were observed.

### Generation of GEMC1-AID mice

Mice were created using CRISPR-Cas9 genome editing. A double stranded DNA donor template targeting the last exon of GEMC1 and containing mAID tagged with mRuby was microinjected along with preassembled crRNA + tracrRNA + Cas9 ribonucleoprotein complexes into B6SJLF1/J mouse embryos at the one-cell stage and transplanted into pseudo pregnant females. Microinjection and transplantation were performed by the Johns Hopkins Transgenic Core Facility. Offspring were screened by PCR and sequencing, and confirmed knock-ins were outcrossed to C57BL6/J mice for two generations. GEMC1-AID-mRuby heterozygous mice were then outcrossed to Rosa26-OsTIRF74G^23^ mice and incrossed to generate homozygous lines.

### Primary mouse tracheal epithelia cell culture

Mouse trachea epithelia cultures were prepared as described previously^24^. Briefly, tracheas were harvested from mice ranging from 2-12 months of age, digested overnight in pronase solution, after which stem cells were harvested and plated on transwell filters. Stem cells were allowed to proliferate for 5-7 days in mTEC Plus media before moving to an air-liquid interface with NuSerum media in the basal chamber to trigger differentiation. The day at which cells were moved to the air-liquid interface was called differentiation day 0. For experiments with DDR kinase inhibitors, verapamil and kinase inhibitor were added to the basal chamber on day 0. DMSO controls were also treated with verapamil. For experiments with etoposide, verapamil was not added. Drugs and concentrations are listed in the Key Resources Table.

### Primary mouse brain ependymal cell culture

Ependymal cells were prepared as described previously^25^. Briefly, brains were harvested from P0-P3 pups, the telecephalons were isolated, enzymatically digested, and progenitor cells were grown to confluence for 4-5 days. Cells were then plated at high density on coverslips and grown in serum-free media to promote differentiation. The day at which cells were moved to serum-free media was called differentiation day 0.

### Immunofluorescence

Cells were fixed in either 4% paraformaldehyde for 15 minutes at room temperature or ice cold 100% methanol for 10 minutes at -20°C. Paraformaldehyde fixed cells were quenched for 5 minutes with glycine. Cells were then washed twice with PBS and permeabilized for 10 minutes in 0.1% Triton-X in PBS. Cells were blocked for 1-2 hours in block buffer (10% normal donkey serum, 0.1% Triton-X in PBS). Primary antibodies were diluted in block buffer and incubated overnight at room temperature. Cells were washed with 0.1% Triton-X in PBS, and then incubated with secondary antibody for 1-2 hours at room temperature. Cells were washed again with 0.1% Triton-X in PBS, and then mounted. Primary and secondary antibodies are listed in the Key Resources Table.

### Confocal microscopy and image analysis

Slides were imaged on either a Zeiss Axio Observer 7 inverted microscope with Slidebook 2023 software (3i—Intelligent, Imaging Innovations, Inc.), CSU-W1 (Yokogawa) T1 Super-Resolution SoRa Spinning Disk, and Prime 95B CMOS camera (Teledyne Photometrics) with a 63x/1.40 NA plan-apochromat oil immersion objective or a Nikon A1R confocal system with Nikon Elements software an 60x/1.40 NA. Images were processed in FIJI^26^. A minimum of three biological replicates were performed for each experiment. All images presented in figures are max projections.

### Comet assay

The alkaline Comet assay^27^ was modified for MCCs. Cells were harvested either on day 0 (stem cells) or day 5 (MCCs) by trypsinization and resuspended in ice cold PBS. 1500 cells were added to 1% agarose in a 1:2 ratio and 100uL of the cell/agarose mixture was quickly pipetted onto an agarose-coated slide. The mixture solidified for 5 minutes and then was submerged in cold lysis solution (1.2M NaCl, 100mM Na2EDTA, 0.1% sodium lauryl sarcosinate, 0.26M NaOH, pH >13) overnight at 4°C. Slides were then submerged in cold rinse solution (0.03M NaOH, 2mM Na2EDTA, pH >12.3) for 1 hour, replacing with fresh rinse solution every 20 minutes. Slides were moved to an electrophoresis chamber with fresh rinse solution just covering the top of the agarose mixture and run at a voltage of 0.6V/cm for 25 minutes. Slides were then neutralized in distilled water followed by staining with 2.5ug/mL propidium iodide for 20 minutes. Cells were rinsed in distilled water and imaged on a Nikon (Ti Eclipse widefield microscope with a 40x/0.75NA air objective and Teledyne Photometrics Kinetix 22 camera. Images were analyzed in FIJI using line scans across the comet to measure intensity values.

### Cell growth and viability assays

To optimize DDR kinase inhibitor concentrations, several assays were performed. First, to measure endpoint growth and viability with prolonged drug treatment, MTT assays were performed as described previously^28^. Briefly, mouse IMCD3 cells were plated in triplicate for each condition, inhibitors were added the following day, and cells were left to proliferate. After 5 days, cell viability was assayed with Thiazolyl blue tetrazolium bromide (MTT) solution and absorbance was measured at 570 nm. Drug titration curves were generated and the highest concentration before cell viability decreased was used for dose response curves in MCCs.

MCCs were treated with a range of drug concentrations surrounding this highest viable concentration from day 0 through day 7, then cells were fixed and analyzed for markers of differentiation and cell viability. The optimal drug concentration was determined by selecting the highest concentration that decreased differentiation but not cell viability/density. The effectiveness of these drug concentrations was validated by loss of immunofluorescence signal when staining with antibodies to the phosphorylated/active form of each DDR kinase.

## Figure Legends

**supplementary Figure 1.**
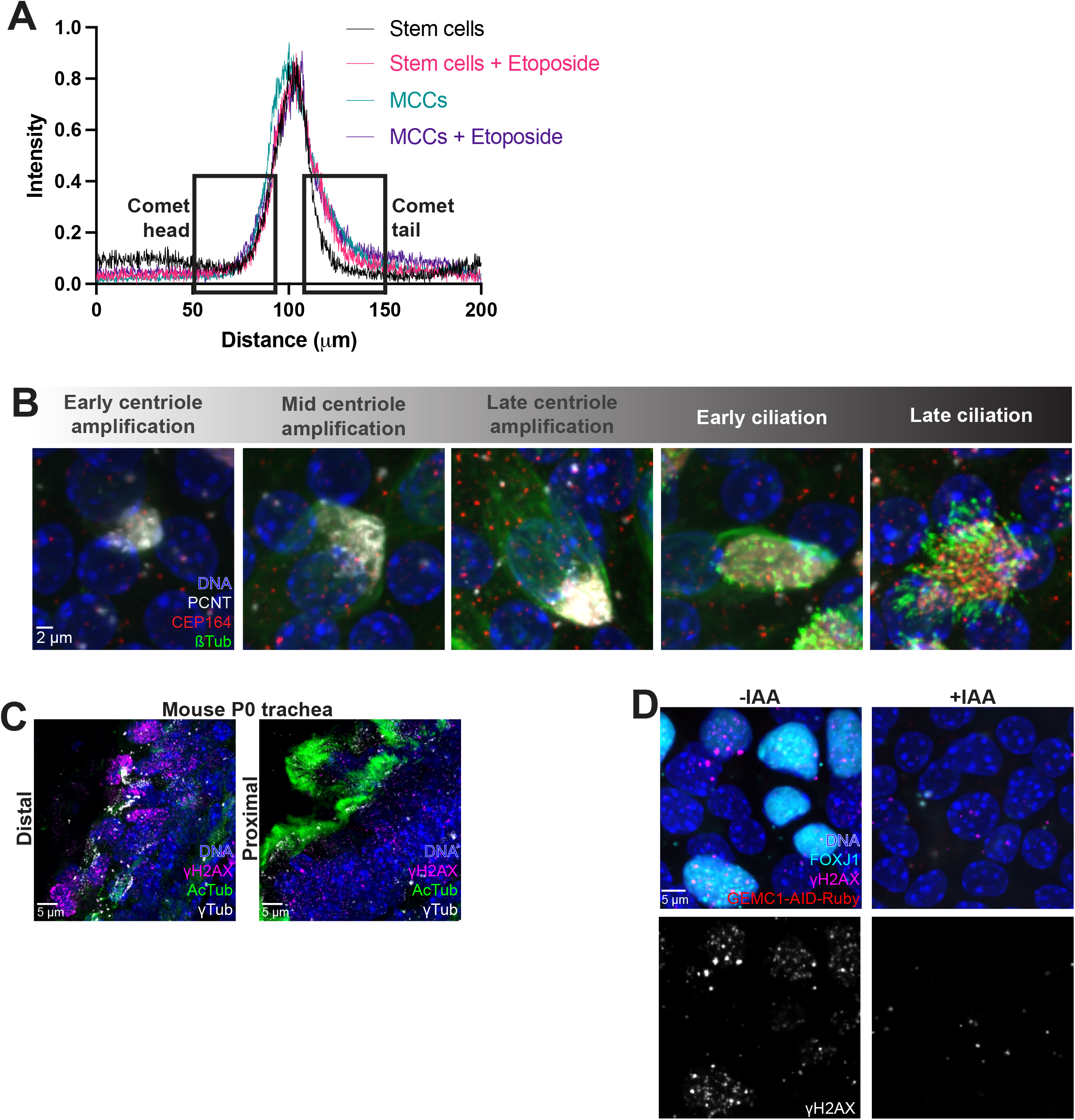
(A) Line scans showing intensity values through the nuclei from representative comet assay images in Figure 1C. The boxed areas were used for quantitation in Figure 1D. (B) (A) Primary mouse trachea epithelia cells at differentiation days 3-7, fixed, and stained for the early centriole protein marker (PCNT, gray), late centriole protein marker (CEP164, red), microtubules, centrioles, and cilia (β-tubulin, green), and DAPI to mark nuclei. Note that cytoplasmic β-tubulin staining denotes mid-late centriole amplification stages. (C) Tissues sections from P0 mouse tracheas fixed and stained for DNA damage (γH2AX, magenta), cilia (AcTub, green), and centrioles (γTub, gray). At P0, MCCs in the trachea are still developing and are more mature proximally than distally. Note the DNA damage occurring in the distal trachea region where cells are undergoing centriole amplification, and less DNA damage in the proximal region where cells are ciliated. (D) Primary mouse trachea epithelia cells at differentiation day 7, fixed, and stained for the MCC transcription factor (FOXJ1, cyan), DNA damage (γH2AX, magenta), and DAPI to mark nuclei. Cells are from GEMC1-AID-Ruby; OsTIR1 mice and were either treated with water (−IAA) or 5-phenyl-1H-indole-3-acetic acid (IAA) from differentiation days 0-7 to degrade GEMC1. While levels of endogenous GEMC1-AID-Ruby protein are below detection by Ruby fluorescence, the loss of MCCs demonstrates effective GEMC1 degradation. Note loss of γH2AX in cells lacking GEMC1.

**supplementary Figure 2.**
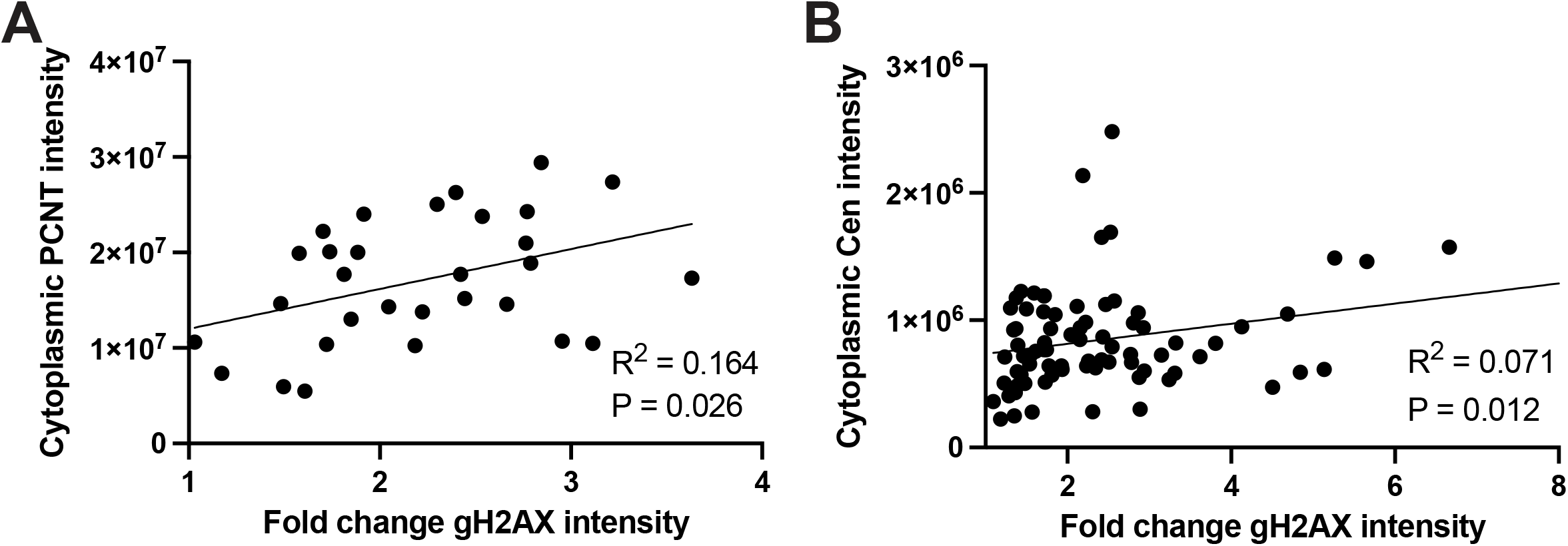
(A-B) Linear correlation plots of cytoplasmic centriole protein intensity (PCNT in (A) or Centrin in (B)) versus fold-change in γH2AX nuclear intensity. Each dot represents a single cell.

**supplementary Figure 3.**
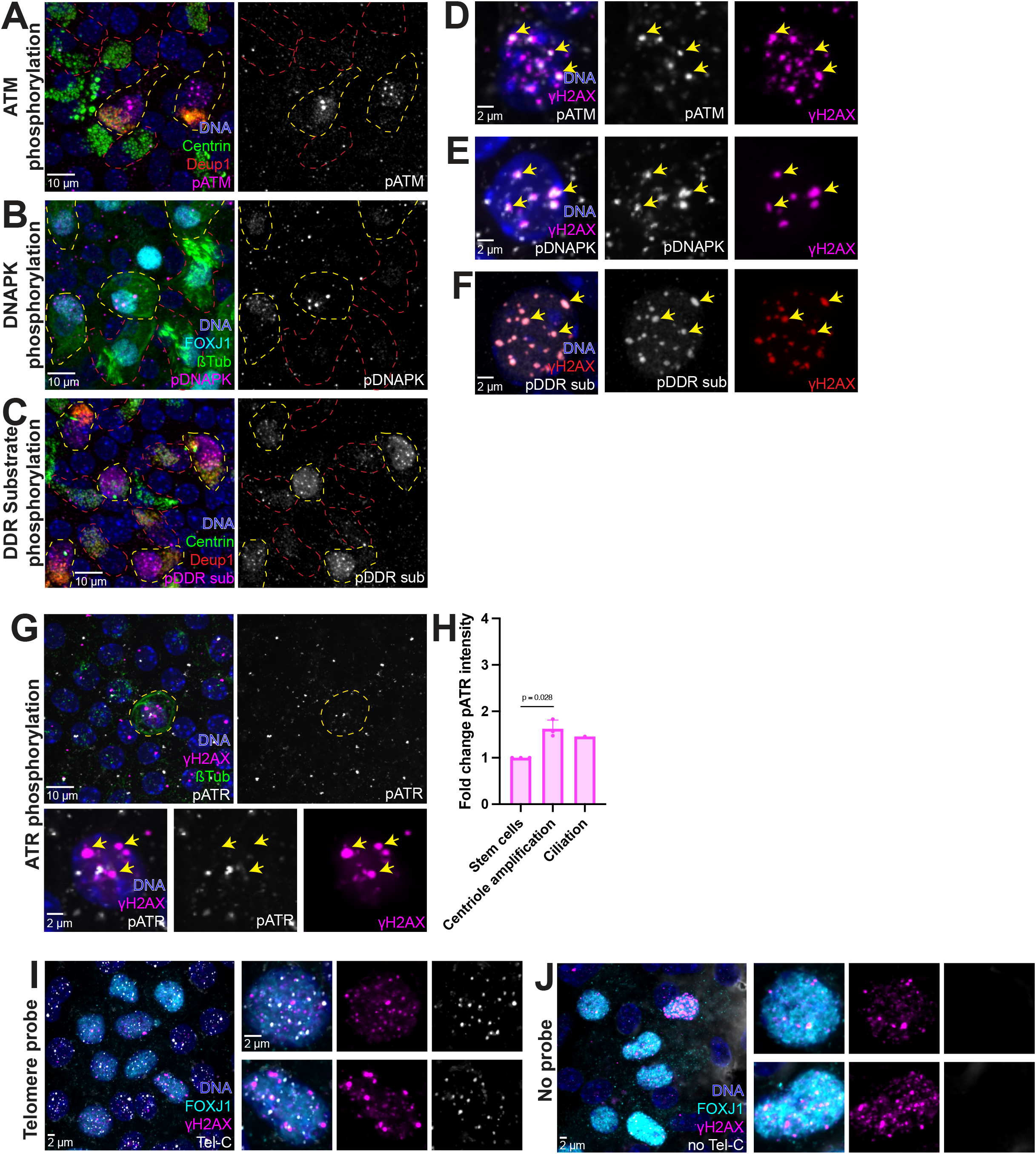
(A) Primary mouse trachea epithelia cells at differentiation day 5, fixed, and stained for centrioles (Centrin, green), deuterosomes (Deup1, red), phosphorylated ATM (pATM, magenta) and DAPI to mark nuclei. MCCs were grouped into differentiation stages based on Centrin and Deup1 staining. Cells outlined in yellow are undergoing centriole amplification and cells outlined in red are at the ciliation stage. (B) Primary mouse trachea epithelia cells at differentiation day 5, fixed, and stained for the MCC transcription factor (FOXJ1, cyan), phosphorylated-DNAPK (pDNAPK, magenta), microtubules, centrioles, and cilia (β-tubulin, green), and DAPI to mark nuclei. MCCs were grouped into differentiation stages based on FOXJ1 and β-tubulin staining. Cells outlined in yellow are undergoing centriole amplification and cells outlined in red are at the ciliation stage. (C) Primary mouse trachea epithelia cells at differentiation day 5, fixed, and stained for centrioles (Centrin, green), deuterosomes (Deup1, red), phosphorylated substrates of the DDR kinases (pDDR sub, magenta) and DAPI to mark nuclei. MCCs were grouped into differentiation stages based on Centrin and Deup1 staining. Cells outlined in yellow are undergoing centriole amplification and cells outlined in red are at the ciliation stage. (D) Primary mouse trachea epithelia cells at differentiation day 5, fixed, and stained for DNA damage (γH2AX, magenta), phosphorylated ATM (pATM, gray) and DAPI to mark nuclei. Arrows denote colocalization between γH2AX and pATM foci. (E) Primary mouse trachea epithelia cells at differentiation day 5, fixed, and stained for DNA damage (γH2AX, magenta), phosphorylated-DNAPK (pDNAPK, gray) and DAPI to mark nuclei. Arrows denote colocalization between γH2AX and pDNAPK foci. (F) Primary mouse trachea epithelia cells at differentiation day 5, fixed, and stained for DNA damage (γH2AX, red), phosphorylated substrates of the DDR kinases (pDDR sub, gray) and DAPI to mark nuclei. Arrows denote colocalization between γH2AX and pDDR foci. (G) Primary mouse trachea epithelia cells at differentiation day 3, fixed, and stained for DNA damage (γH2AX, magenta), microtubules, centrioles, and cilia (β-tubulin, green), phosphorylated ATR (pATR, gray), and DAPI to mark nuclei. The cell outlined in yellow is undergoing centriole amplification and yellow arrows in bottom panels denote lack of colocalization between γH2AX and pATR foci. (H) Fold-change in pATR nuclear intensity compared to neighboring stem cells. (I-J) DNA-FISH with the Tel-C telomere probe (I) or no probe (J) combined with immunofluorescence with the MCC transcription factor (FOXJ1, cyan), DNA damage (γH2AX, magenta), and DAPI to mark nuclei. Note the lack of colocalization between γH2AX foci and telomeres.

**supplementary Figure 4.**
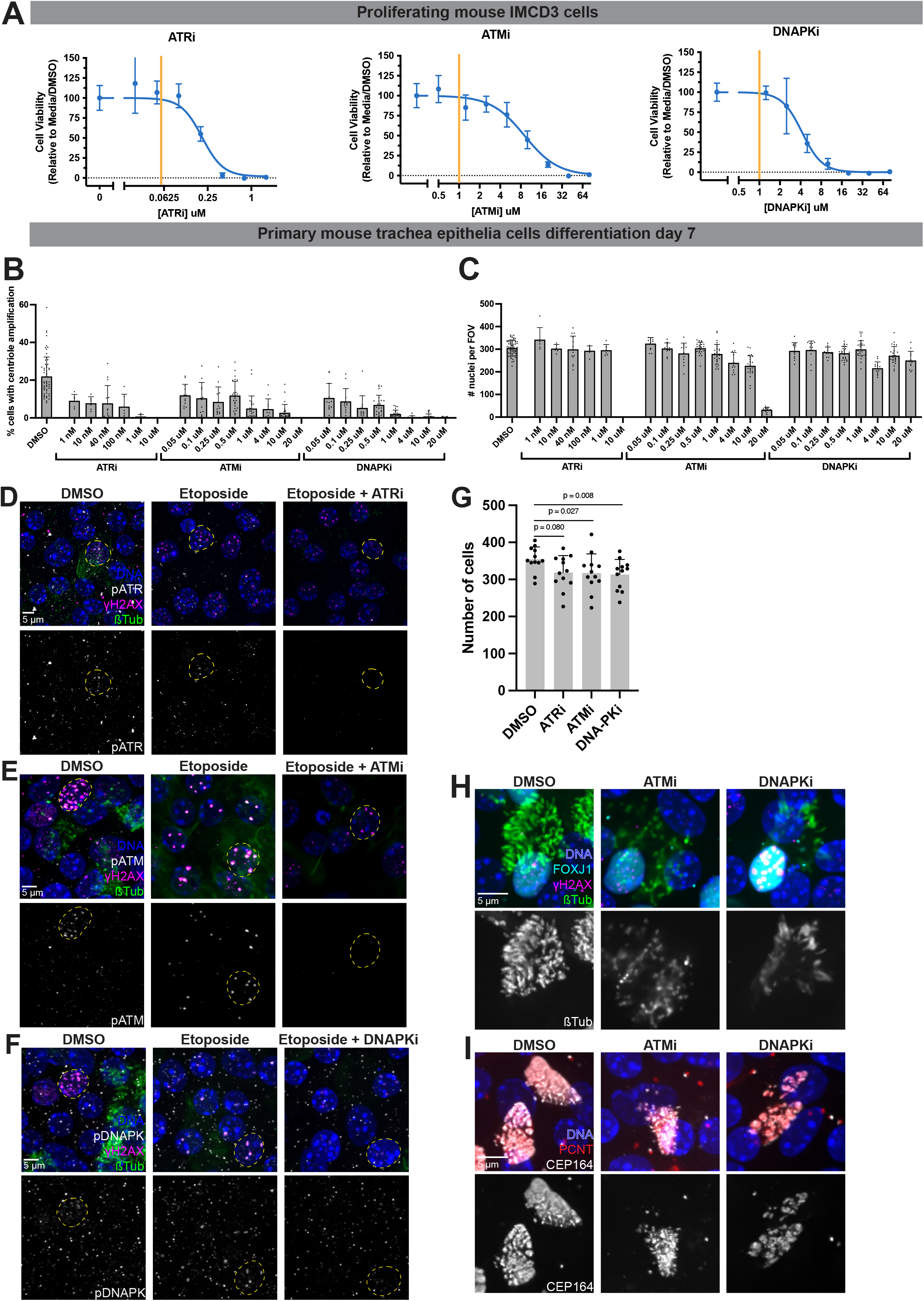
(A) MTT endpoint viability assay for titration of DDR kinase drugs in proliferating mouse IMCD3 cells. Yellow line denotes concentrations used for experiments. (B-C) Dose response curves of DDR kinase drugs in primary mouse trachea epithelia cells treated from differentiation days 0-7, fixed, and stained for markers of centriole amplification (B) and DAPI to determine number of cells (C). (D) Primary mouse trachea epithelia cells at differentiation day 3 fixed, and stained for DNA damage (γH2AX, magenta), phosphorylated ATR (pATR, gray), microtubules, centrioles, and cilia (β-tubulin, green), and DAPI to mark nuclei. Cells were treated with either DMSO, etoposide, or etoposide plus ATRi to validate phospho-antibody and kinase inhibitor treatment. Cells outlined in yellow show pATR foci that are lost with ATRi treatment. (E) Primary mouse trachea epithelia cells at differentiation day 5, fixed, and stained for DNA damage (γH2AX, magenta), phosphorylated ATM (pATM, gray), microtubules, centrioles, and cilia (β-tubulin, green), and DAPI to mark nuclei. Cells were treated with either DMSO, etoposide, or etoposide plus ATMi to validate phospho-antibody and kinase inhibitor treatment. Cells outlined in yellow show pATM foci that are lost with ATMi treatment. (F) Primary mouse trachea epithelia cells at differentiation day 5, fixed, and stained for DNA damage (γH2AX, magenta), phosphorylated DNAPK (pDNAPK, gray), microtubules, centrioles, and cilia (β-tubulin, green), and DAPI to mark nuclei. Cells were treated with either DMSO, etoposide, or etoposide plus DNAPKi to validate phospho-antibody and kinase inhibitor treatment. Cells outlined in yellow show pDNAPK foci that are lost with DNAPKi treatment. (G) Graph showing the number of cells with either DMSO or kinase inhibitor treatment from Figure 4A and B. (H) Primary mouse trachea epithelia cells at differentiation day 7, fixed, and stained for the MCC transcription factor (FOXJ1, cyan), DNA damage (γH2AX, magenta), microtubules, centrioles, and cilia (β-tubulin, green), and DAPI to mark nuclei. Cells were treated with either DMSO or indicated kinase inhibitor. Note the differences in cilia (β-tubulin staining) between DMSO and kinase inhibitor conditions. (I) Primary mouse trachea epithelia cells at differentiation day 5, fixed, and stained for early centriole protein marker (PCNT, gray), late centriole protein marker (CEP164, red), microtubules, centrioles, and cilia (β-tubulin, green), and DAPI to mark nuclei. Cells were treated with either DMSO or indicated kinase inhibitor. Note the differences in centrioles (CEP164) between DMSO and kinase inhibitor conditions. All graphs show mean ± SD.

